# A process-based metacommunity framework linking local and regional scale community ecology

**DOI:** 10.1101/832170

**Authors:** Patrick L. Thompson, Laura Melissa Guzman, Luc De Meester, Zsófia Horváth, Robert Ptacnik, Bram Vanschoenwinkel, Duarte S. Viana, Jonathan M. Chase

## Abstract

The metacommunity concept has the potential to integrate local and regional dynamics within a general community ecology framework. To this end, the concept must move beyond the discrete archetypes that have largely defined it (e.g. neutral vs. species sorting) and better incorporate local scale species interactions and coexistence mechanisms. Here, we present a fundamental reconception of the framework that explicitly links local coexistence theory to the spatial processes inherent to metacommunity theory, allowing for a continuous range of competitive community dynamics. These dynamics emerge from the three underlying processes that shape ecological communities: 1) density-independent responses to abiotic conditions, 2) density-dependent biotic interactions, and 3) dispersal. Stochasticity is incorporated in the demographic realization of each of these processes. We formalize this framework using a simulation model that explores a wide range of competitive metacommunity dynamics by varying the strength of the underlying processes. Using this model and framework, we show how existing theories, including the traditional metacommunity archetypes, are linked by this common set of processes. We then use the model to generate new hypotheses about how the three processes combine to interactively shape diversity, functioning, and stability within metacommunities.

**Statement of authorship:** This project was conceived at the sTURN working group, of which all authors are members. PLT developed the framework and model with input from all authors. PLT wrote the model code. PLT and LMG performed the simulations. PLT produced the figures and wrote the first draft with input from LMG and JMC. All authors provided feedback and edits on several versions of the manuscript.

**Data accessibility:** All code for running the simulation model and producing the figures is archived on Zenodo - https://doi.org/10.5281/zenodo.3833035.

## Introduction

The field of community ecology encompasses a large number of theories, concepts, and hypotheses (Mittelbach & McGill 2019). These include the ecological niche (Grinnell 1917; Elton 1927; Hutchinson 1957), the resource ratio hypothesis (Tilman 1982), neutral theory (Bell 2000; Hubbell 2001), coexistence theory (Chesson 2000b), and the relationship between coexistence-mediated biodiversity and the functioning of ecosystems (Tilman 1999; Loreau & Hector 2001). These concepts, however, are far from unified, and it is often unclear how they relate to one another, or whether they are compatible (e.g. niche versus neutral-based perspectives). This is because theories often differ in their assumptions, the spatial and temporal scales to which they apply, the notation used to express them mathematically, and the degree to which they are phenomenological versus mechanistic.

Following a period of time where the multitude of conceptual frameworks and empirical approaches led to community ecology being labelled “a mess” (Lawton 1999), more recent efforts have led towards reconciliation. For example, Gravel et al. (2006) demonstrated how niche-like and neutral-like dynamics are opposite ends of a continuum defined by the degree to which species overlap in their abiotic niches (see also Adler *et al*. 2007). More generally, Vellend (2010; 2016) categorized community ecological processes into broad groupings (e.g., selection, drift, dispersal, and speciation) in an attempt at synthesis. Nevertheless, a key divide in community ecology remains between theories that focus on the dynamics of species interactions and coexistence within local habitats versus those that focus on the dynamics of regional metacommunities that form across heterogeneous habitats and are connected by dispersal.

The metacommunity concept has the potential to serve as a unifying framework for community ecology theory. The concept posits that metacommunities are composed of sets of local habitats that are connected by dispersal, and that species within each local habitat interact with each other and respond to local environmental conditions (Leibold & Chase 2018). Thus, the framework has the potential to incorporate theories that focus on the dynamics of local habitats as well as theories that involve regional connectivity and environmental heterogeneity across space.

Still, two main aspects of the metacommunity concept have limited its uptake as a general framework for understanding the underlying pattern and process in community ecology. First, since its initial synthesis (Leibold *et al*. 2004), metacommunity theory has most often been expressed using discrete and seemingly incompatible archetypes (i.e. neutral dynamics, patch dynamics, species sorting, and mass effects) rather than through a series of continuous processes that are common to all communities (Logue *et al*. 2011; Brown *et al*. 2017). More recent models are capable of generating the metacommunity archetypes by altering key parameters (e.g. dispersal, niche breadth, species interactions, and stochasticity), bringing us closer to the goal of redefining metacommunity theory based on processes (e.g. Shoemaker & Melbourne 2016; Fournier *et al*. 2017; Ovaskainen *et al*. 2019). Yet, these models only consider discrete ranges of parameter space corresponding to the archetypes, and so remain focused on the archetypes rather than on the underlying processes. Second, most metacommunity theory oversimplifies local scale dynamics by assuming, either implicitly or explicitly, that all species compete equally (e.g. Hubbell 2001; Loreau *et al*. 2003; Shoemaker & Melbourne 2016; Worm & Tittensor 2018). In these models, diversity is maintained by spatial coexistence mechanisms (Chesson 2000a), while local coexistence mechanisms tend to be missing (Shoemaker & Melbourne 2016).

In contrast, biotic interactions, in particular competition for resources, are central to theories about local scale coexistence. The idea that species must compete more strongly with themselves than with each other to coexist dates back to Gause’s competitive exclusion principle (Gause 1932), based on Lotka-Volterra models (Lotka 1922; Volterra 1926), and is the mainstay of contemporary theories of coexistence (e.g. Tilman 1982; Chesson 2000b; Chase & Leibold 2003). When these conditions are not met, other local dynamics can emerge, including neutral dynamics and priority effects (Cushing *et al*. 2004; Adler *et al*. 2007; Ke & Letten 2018). Modern coexistence theory (Chesson 2000b) includes spatial coexistence mechanisms that align with those of metacommunity theory (Shoemaker & Melbourne 2016). However, applications of coexistence theory rarely account for dispersal or spatial environmental heterogeneity.

Here, we propose a general process-based metacommunity framework that unites local and regional scale theory of ecological community dynamics. We do this by reframing metacommunity theory and local coexistence theory based on three common core processes that govern the dynamics of communities and that are rooted in classic theory. These are: 1) density-independent responses to abiotic conditions, 2) density-dependent biotic interactions, and 3) dispersal, as well as stochasticity in how these processes impact demography. Our processes differ slightly from Vellend’s (2010; 2016) categorizations for a few reasons. First, we ignore the processes of speciation for the purposes of tractability, although in our discussion we propose how evolution of the traits that underly these processes could be considered as a fourth process. Second, we separate selection-based processes depending on whether their effects on populations depend on density, which is critical because density-independent and dependent processes affect coexistence in fundamentally different ways (Chase & Leibold 2003), and, as we outline below, they correspond to different parameters in classic ecological theory. Third, while stochasticity is a critical feature of our framework, we do not consider it as a separate process but instead consider stochasticity as an inherent feature in the realization of the three fundamental processes—abiotic responses, biotic interactions, and dispersal—which ultimately generates demographic stochasticity (Shoemaker *et al*. 2019). By distinguishing processes in this way, we can unite metacommunity theory and local coexistence within a single mathematical framework. Furthermore, we can use this more generalized theory to make explicit predictions about a number of patterns including the influence of varying dispersal rates on patterns of local and regional diversity, and on the relationship between local biodiversity and the functioning of the ecosystems.

### A general framework for competitive metacommunities

We formalize a metacommunity as a set of local communities where populations of multiple species potentially compete and can disperse among local communities that are distributed in space. Figure 1 depicts a schematic of such a metacommunity, visualizing a landscape in which local communities are separated via an uninhabitable matrix and connected via dispersal. Nevertheless, our modelling framework need not apply to only this canonical view of a metacommunity and, in fact, all communities exist within a larger landscape (i.e., metacommunity) characterized by differences in scale (i.e. local and regional), heterogeneity, and dispersal (Leibold & Chase 2018). We provide an overview of four basic propositions that form the foundation for a generalized metacommunity ecology framework.

**Figure 1.**
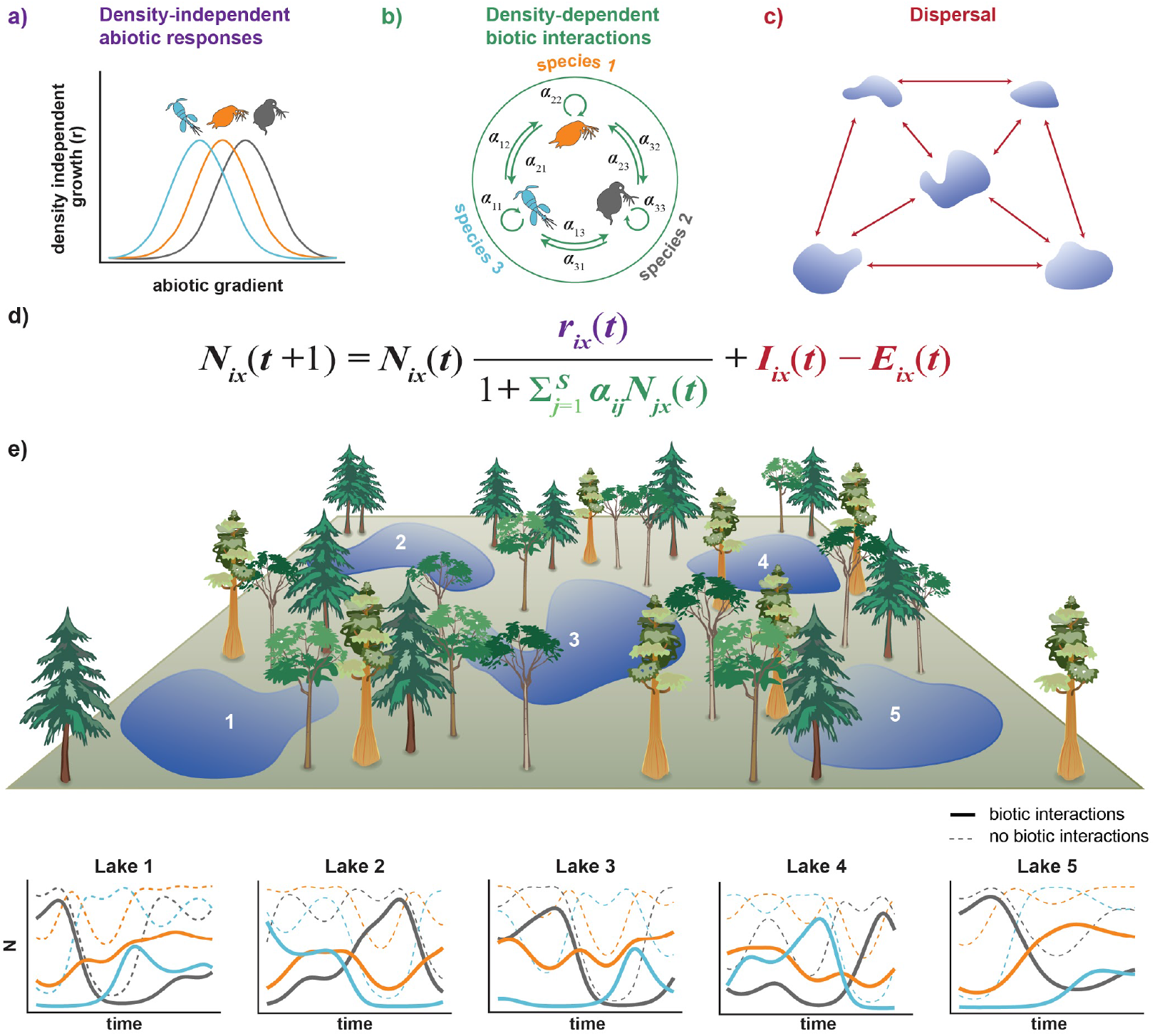
A schematic representation of our metacommunity framework and how we formalize each aspect of it in our mathematical model. a) Density-independent abiotic niches of three zooplankton species are represented graphically, where *r_i_* follows a Gaussian response curve over the gradient of abiotic environmental conditions in the metacommunity, but each species *i* has a different environmental optimum. b) Dynamics also depend on interactions within and among species. This is included as *per capita* intraspecific *α_ii_* interspecific *α_ij_* interaction coefficients, and their realized impact on population dynamics increases with population size *N_ix_*(*t*). Note, *α_ij_* could be made to vary with environmental conditions, but in this paper we have assumed that it does not. c) Dispersal alters population sizes via immigration and emigration and depends on the physical arrangement of habitat patches in the landscape. d) Each of these processes is expressed as separate expressions in our mathematical model. In this model, *N_ix_*(*t*) is the abundance of species *i* in patch *x* at time *t*, *r_ix_*(*t*) is its density-independent growth rate, is the *per capita* effect of species *j* on species *i, I_ix_*(*t*) is the number of individuals that arrive from elsewhere in the metacommunity via immigration, and *E_ix_*(*t*) is the number of individuals that leave via emigration. e) Simulated dynamics of a three zooplankton species, five lake metacommunity. The abiotic conditions vary across time and space and species respond to this heterogeneity via the Gaussian response curves in panel a. The species also compete so that the realized dynamics differ from those that would occur in the absence of interspecific competition (thick solid vs. thin dashed lines). Dispersal connects populations via immigration and emigration, with more individuals being exchanged between lakes that are in close proximity. Although stochasticity in population growth and dispersal is integral to our framework and is included in all other simulations presented in this paper, we have omitted stochasticity from these dynamics to increase clarity. Figure design by Sylvia Heredia.

#### 1) *The density-independent growth rate of a population depends on the local abiotic conditions* (Figure 1a)

Density-independent growth in the absence of intra- or interspecific competition is determined by the dimensions of the abiotic environment (e.g. temperature and rainfall) that influence organism performance but where the organisms do not impact that abiotic dimension (Tilman 1982; Chase & Leibold 2003). Thus, density-independent growth defines the range of conditions that allow for positive intrinsic growth (i.e. fundamental niche; Hutchinson 1957). In classic models (e.g. Lotka-Volterra), density-independent growth is expressed using the constant *r*.

Density-independent abiotic conditions vary in space and time, resulting in differences in density-independent growth (*r*), depending on the shape of the species’ abiotic niche. Abiotic niches are often thought to have a Gaussian shape, but they can take any form, including skewed (e.g. thermal performance curves) or positive (e.g. growth over increasing nutrient supply). When abiotic niche curves are narrow over the range of abiotic conditions, species will respond strongly to this variation (closer to the assumptions of classical ‘niche-based’ theory). In contrast, if abiotic niches are broad or flat over the range of abiotic conditions experienced, species will respond weakly or not at all (closer to the assumptions of neutral theory).

Density-independent responses to environmental conditions have been the focus of most niche-based metacommunity models (e.g. Loreau *et al*. 2003; Gravel *et al*. 2006; Shoemaker & Melbourne 2016; Fournier *et al*. 2017). However, density-independent responses to the environment are only one aspect of the realized niche, and this has led to confusion in how niche processes govern the dynamics of metacommunities. Coexistence within a habitat patch is more likely when species have similar density-independent growth rates, because it makes equal fitness more likely (sensu Chesson 2000b; Godoy & Levine 2014). But, local scale coexistence also requires stabilizing density-dependent biotic interactions or dispersal (Chesson 2000b; Snyder & Chesson 2004).

#### 2) *The realized growth rate of a population depends on the density-dependent intra- and interspecific interactions* (Figure 1b)

Density-dependent competition limits species growth and, along with density-independent abiotic responses, determines equilibrium abundances (i.e. carrying capacity Verhulst 1838; Lotka 1922; Volterra 1926). Density-dependent aspects of the environment can include abiotic variables, such as resource availability, which are altered by the populations that consume them. Likewise, density-dependent aspects of the environment can include the population size of other organisms, such as predators, prey, or mutualists. In all cases, the impacts of density-dependent aspects of the environment can be expressed as *per capita* interactions. As our focus here is on competitive communities, we assume that biotic interactions represent competition for common resources but recognize that other interaction types can be included in these models in a relatively straightforward way.

Most niche-based metacommunity theory assumes that species do not differ in their resource use (i.e. equal intra- and interspecific competition), but rather their competitive ability depends only on their match with the density independent aspects of the local environment (e.g. Loreau *et al*. 2003; Gravel *et al*. 2006; Shoemaker & Melbourne 2016; Fournier *et al*. 2017; Worm & Tittensor 2018; but see Liautaud *et al*. 2019). Yet, we know from local coexistence theory and empirical evidence that this is rarely the case (Chesson 2000b; Adler *et al*. 2018). Density-dependent competition results because organisms impact resource availability (e.g. space, nutrients, water availability, prey). The density-dependent interaction strength of any species with its competitors for shared resources depends on the degree of overlap in resource use (sometimes termed ‘niche differentiation’; Chesson 2000b). In classic Lotka-Volterra models, density-dependent interactions are expressed using the *α_i,j_*, which is the *per capita* impact of species *j* on species *i*. *α_i,j_* is a phenomenological parameter and is just one way that density-dependent biotic interactions can be expressed (Tilman 1982; Letten *et al*. 2017) but is useful to understand how biotic interactions can lead to a range of dynamics.

The density-dependent aspect of the niche has very different implications for community dynamics compared to the density-independent abiotic niche responses outlined above. In a community of two species that have equal intraspecific competition (i.e. *α_i,i_* = *α_j,j_*), equal density-independent growth (i.e. *r_i_* = *r_j_*), and thus equal equilibrium abundance, four different outcomes are possible within a single habitat, depending on the balance of inter- (i.e. *α_i,j_*) to intraspecific (i.e. *α_i,i_*) competition (Cushing *et al*. 2004):

a. equal competition - individuals of all species compete equally (i.e. *α_j,i_* = *α_i,i_* = *α_i,j_* = *α_j,j_*), so that coexistence is unstable but exclusion is slow and determined by stochastic processes;
b. stabilizing competition - species compete more strongly with themselves than with each other (i.e. *α_j,i_* < *α_i,i_* and *α_i,j_* < *α_jj_*);
c. competitive dominance - competition is unbalanced (i.e. *α_j,i_* > *α_i,i_* and *α_i,j_* < *α_j,j_*) so that the superior competitor *i* excludes the inferior species *j*, regardless of initial abundances;
d. destabilizing competition - species compete with each other more strongly than with themselves (i.e. *α_j,i_* > *α_i,i_* and *α_i,j_* > *α_j,j_*) so that the species with higher initial abundance excludes the other species.

Of course, the outcome of competition becomes more complicated in more diverse communities (Barabás *et al*. 2016; Saavedra *et al*. 2017), when species have different strengths of intraspecific competition and have different density-independent growth rates (Godoy & Levine 2014). Nevertheless, these four outcomes set the context for the dynamics of multispecies communities by determining whether local scale communities have single or multiple attractors.

Because density-dependent biotic interactions are strong determinants of local scale dynamics, they also influence how species respond to environmental change. This is because density-independent abiotic environmental factors can alter the abundance of interacting species, which effectively modifies density-dependent interactions (Ives & Cardinale 2004). These indirect effects of the environment are often strong in empirical communities (Davis *et al*. 1998; Alexander *et al*. 2015) but are absent from models assuming that species all interact equally. By including density-dependent biotic interactions, we can explore how the range of community dynamics that result from biotic interactions at local scales interact with spatial metacommunity processes.

#### 3) *The size of a population depends on dispersal* (Figure 1c)

Dispersal modifies the dynamics of local populations and communities both directly and indirectly. Emigration reduces population size, while immigration increases population size and can bring in species that, through their density-dependent biotic interactions, strongly impact community structure. Furthermore, dispersal provides additional mechanisms that can allow species to coexist at local scales (Chesson 2000b; Snyder & Chesson 2004).

Along a gradient from low to high dispersal, we expect to see the following processes (Mouquet & Loreau 2003):

a. low dispersal rates result in dispersal limitation, where spatial isolation prevents potential colonizers;
b. intermediate dispersal allows species to colonize new habitats, allowing for environmental tracking, rescue effects, and recolonization;
c. high dispersal rates may facilitate source-sink dynamics whereby immigration increases population size in locations where local conditions are less favourable and emigration decreases population size in locations where conditions are more favourable, thus promoting the spatial homogenization of metacommunities.

Multiple dispersal-mediated processes are likely to occur simultaneously within a metacommunity, especially when patches are not equally connected (Thompson *et al*. 2017).

Dispersal is a spatially-explicit process that depends on the spatial connectivity of landscapes, the distance between habitats, species dispersal traits (e.g. dispersal rate and kernel), and population sizes. Metacommunity models have typically been spatially implicit for tractability, but explicit space can be an important determinant of dynamics (e.g. Fournier *et al*. 2017; Thompson *et al*. 2017).

Because dispersal alters population sizes, it also alters the realized strength of density-dependent processes and competition (Holt 1985). Likewise, population sizes alter the number of dispersing individuals, creating a feedback between dispersal and biotic interactions. This contrasts with the view that dispersal, the abiotic environment, and the biotic environment form a series of hierarchical filters to determine community assembly (e.g. Vellend 2016).

#### 4) Births, deaths, immigration, and emigration are stochastic processes that prevent metacommunity dynamics from being purely deterministic

Stochasticity is inherent to any ecological system (McShea & Brandon 2010; Vellend 2016). Yet, rather than incorporating it as a separate process *per se* (e.g. Vellend 2016), we consider it to be an element of each of the three processes. This is because biological processes are probabilistic, resulting in stochasticity in demography and dispersal (Shoemaker *et al*. 2019), which aligns with the long history of models that have included it via probabilistic draws of the underlying biological processes (e.g. Levins & Culver 1971; Hubbell 2001; Matias *et al*. 2012; Shoemaker & Melbourne 2016; Fournier *et al*. 2017). Stochasticity in environmental conditions then alters metacommunity dynamics via the three processes we emphasize here (Shoemaker *et al*. 2019).

### Delineating the range of possible competitive metacommunity dynamics

We illustrate this framework of a continuum of processes in three-dimensional space, with the axes referring to density-independent abiotic responses, density-dependent biotic interactions, and dispersal (Figure 2). While such an illustration is imperfect because these processes are not strictly one dimensional (e.g. dispersal depends on both rate and distance), it is a useful heuristic that allows us to compare and relate different metacommunity dynamics across this three-dimensional parameter space. The first axis is defined by the strength of density-independent abiotic responses. At one extreme, species have no response to heterogeneity in density independent abiotic variables (i.e. flat abiotic niches) and at the other extreme, heterogeneity results in large differences in density-independent growth (i.e. narrow abiotic niches). The second axis is defined by the degree to which competition for resources is stabilizing (i.e. *α_i,j_* < *α_i,i_*), equal (i.e. *α_i,j_* = *α_i,i_*), or destabilizing (i.e. *α_i,j_* = *α_i,i_*) for a given pair of species. The final axis is defined by the probability of dispersal, ranging from no dispersal to the limit where all individuals disperse in every time step. Within a given metacommunity, it is possible that species will differ in their positioning on these axes, for example, if species have different dispersal rates, abiotic niche breadths, and competitive strengths.

**Figure 2.**
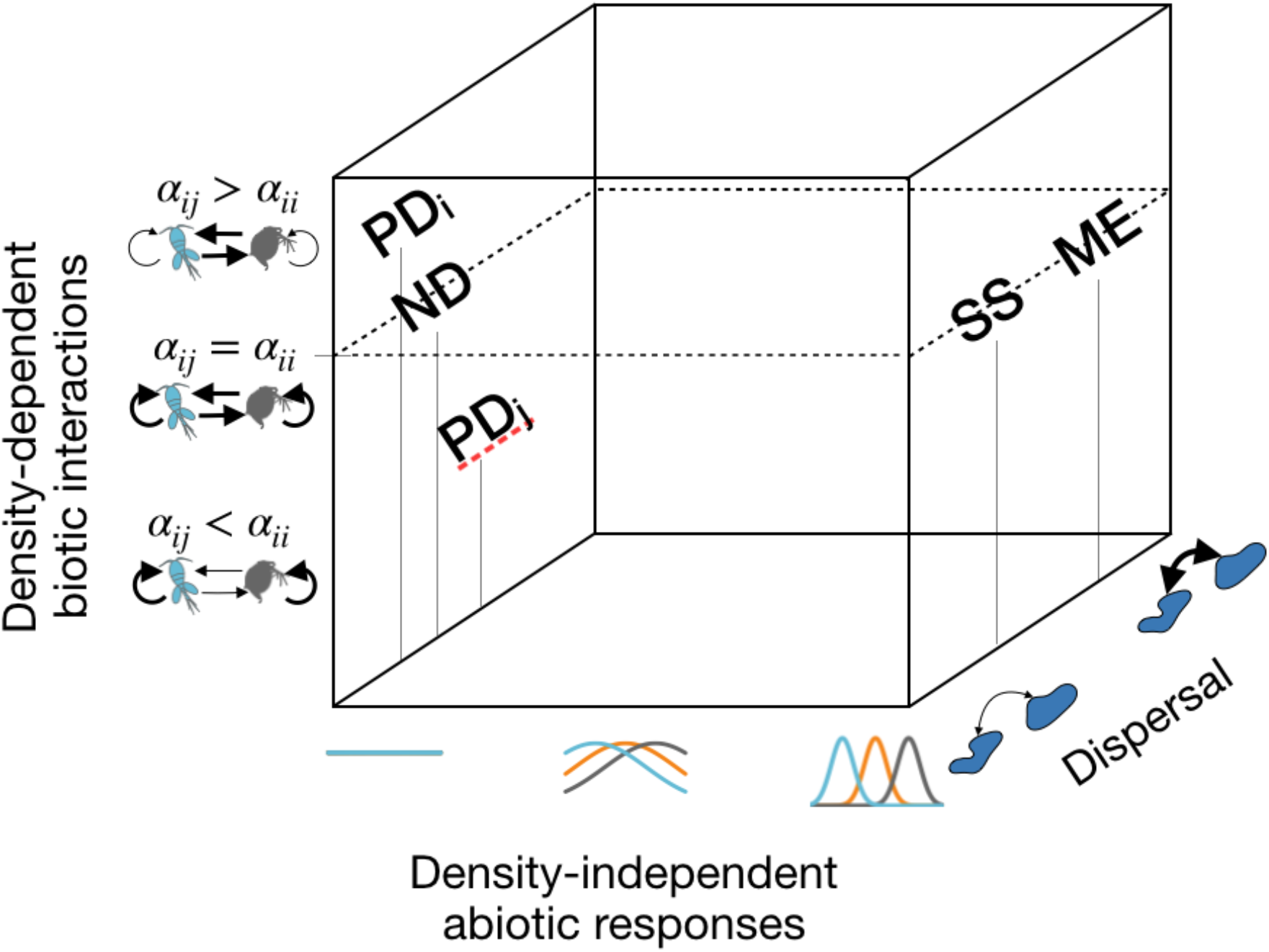
Illustration of the three dimensions of possible metacommunity dynamics. Three key dimensions of our framework define this space: 1) density-independent abiotic responses that range from a flat abiotic niche to narrow abiotic niches with interspecific variation in optima, 2) density-dependent biotic interactions that range depending on the relative strength of interspecific and intraspecific interactions, and 3) dispersal that ranges from very low dispersal rates to very high dispersal rates. The approximate location of each of the four original metacommunity archetypes: ND - neutral dynamics, PD - patch dynamics, SS - species sorting, and ME - mass effects, is indicated to illustrate how our framework links to previous theory. The lines below each label indicate their position in x,z space. PD_i_ indicates the position for competitively dominant species with lower dispersal, PDj indicates the position for competitively weaker species with higher dispersal. Importantly, much of this space is undefined by the four archetypes but represents potential dynamics that can emerge from different combinations of the three processes of our framework.

The traditional organization of metacommunity theories into archetypes (Leibold *et al*. 2004) is consistent with this framework (Box 1). In articulating them based on the three defining processes, we can see how they relate to one another (Figure 2). It also illustrates that these traditional archetypes encompass only a small subset of the possible parameter space of dynamics (Figure 2); other theories have explored some of the space between the archetypes (e.g. Leibold & Chase 2018), but it is clear that there is much more conceptual space to explore.

#### Box 1 the metacommunity archetypes expressed using the three processes

##### Neutral dynamics

The neutral archetype assumes that species have equal density-independent responses to abiotic heterogeneity, that density-dependent competition between species is equal, and that dispersal limitation is acting (i.e. dispersal is not high enough to homogenize the metacommunity) (Figure 2). Equal density-independent responses can be the result of indifference to environmental heterogeneity (i.e. flat abiotic niches) (e.g. Hubbell 2001) or of species having identical responses to abiotic heterogeneity (e.g. Gravel *et al*. 2006). Critically, this archetype implicitly assumes that competition between species is equal. Because of these particularly limiting assumptions, neutral dynamics occur only in a very confined region of our metacommunity parameter space.

##### Species sorting and mass effects

The species sorting archetype assumes that density-independent responses to abiotic heterogeneity are strong and that they differ amongst species (Figure 2). It makes no explicit assumptions about density-dependent biotic interactions, although many models assume equal inter- and intraspecific competition (but see Chase & Leibold 2003; Thompson & Gonzalez 2017). Finally, the archetype implicitly assumes that dispersal is sufficient so that species can access favourable habitat patches, but that dispersal is not so high as to homogenize the metacommunity. The mass effects archetype makes these same assumptions, but, in this case, dispersal rates are higher, allowing populations to persist in habitats that are otherwise unsuitable for growth (Figure 2).

##### Patch dynamics

The patch dynamics archetype has more complicated assumptions, including interspecific variation in competitive ability (e.g., stronger competitors exclude weaker competitors when they are both present in the same habitat patch). Classic models were agnostic to whether these competitive differences were due to density-dependent or density-independent processes (Levins & Culver 1971; Hastings 1980; but see Tilman 1994). More recently, models have included the assumption of competitive differences in density-independent responses to the abiotic environment (i.e. competitively dominant species have higher density-independent growth), while assuming that density-dependent biotic interactions are equal (e.g. Shoemaker & Melbourne 2016; Fournier *et al*. 2017). Nevertheless, it is equally possible that such competitive exclusion can arise from unbalanced density-dependent competition (i.e. *α_i,j_* > *α_j,i_* and *α_j,i_* < *α_j,j_*), with equal density-independent abiotic responses. This is how we have modelled the competition-colonization trade-off here (see Figure 2). For species to coexist in the patch dynamics archetype, we must also assume that there is a competition-colonization trade-off; species persist regionally because competitively weaker species have a higher probability of dispersing. Patch dynamics also implicitly assumes the occurrence of periodic disturbances, be they stochastic extinctions or environmental perturbations, that cause local extirpations.

### Model description

The framework discussed above is quite general and can accommodate a number of different formalizations. For our purposes here, we formalized these assumptions into a simulation model using Beverton-Holt discrete time logistic population growth with Lotka-Volterra competition and spatially-explicit dispersal. The model simulates abundance-based population dynamics of *S* interacting species in *M* habitat patches, coupled by dispersal (Figure 1d, e). Thus, after accounting for density-independent responses to the abiotic environment, densitydependent competition (following Beverton & Holt 1957), dispersal, as well as stochasticity in these responses, *N_ix_*(*t* + 1) is the population size of species *i* in patch *x* at time *t* + 1:

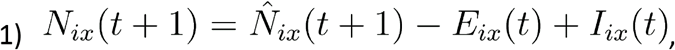

where 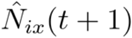 is the population size at time *t* + 1, before accounting for dispersal. 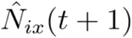 is determined by:

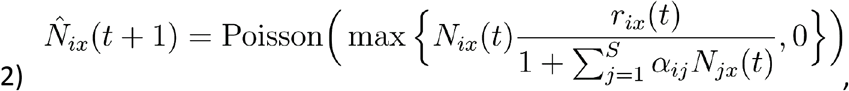

where *N_ix_*(*t*) is the population size at time *t*, and accounts for density-independent abiotic responses *r_ix_*(*t*) and density-dependent biotic interactions *α_ij_* as outlined below. The Poisson distribution ensures integer population sizes and incorporates demographic stochasticity, as outlined below. Note, that a value of zero is assigned to 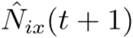 for any Poisson draw with a λ of 0.

#### Density-independent growth

We have assumed that the density-independent abiotic environment is defined by a single variable that varies across space and time, and that *r_ix_*(*t*) is a Gaussian function of this environmental gradient such that

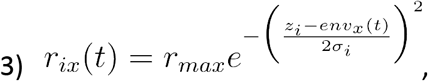

where *r_max_* is the maximum density-independent growth rate, *z_i_* is the environmental optimum of species *env_x_*(*t*) is the environmental conditions in patch *x* at time *t*, and *σ_i_* is the abiotic niche breadth, which determines the rate at which growth is reduced by a mismatch between *z_i_* and *env_x_*(*t*) We manipulate the strength of density-independent responses to abiotic heterogeneity through the parameter *σ_i_*. When *σ_i_* is small, the abiotic niche is narrow and so performance drops quickly when species are present in suboptimal environmental conditions. In contrast, when *σ_i_* is large, the abiotic niche is broad. As *σ_i_* becomes larger relative to the range of environmental conditions in the landscape, the abiotic niche effectively becomes flat, so that species growth is unaffected by environmental heterogeneity.

#### Density-dependent biotic interactions

Density-dependent biotic interactions are modelled using *per capita* interaction coefficients *α_ij_* as outlined in the framework section above. We generate different competitive scenarios by manipulating the relative strength of interspecific (*α_ij_*) and intraspecific (*α_ii_*) competition. We assume that these interaction coefficients are fixed across time and space—in effect assuming that *per capita* interactions do not depend on density-independent aspects of the environment (Tilman 1982). In reality, however, we know that *per capita* competition strengths can vary between habitats and with environmental conditions (Germain *et al*. 2018; Wainwright *et al*. 2018; Grainger *et al*. 2019). Allowing *per capita* interaction strengths to vary with environmental conditions is an important next step but would complicate the model substantially and is not necessary to meet the goals of this paper.

#### Dispersal

Dispersal is incorporated via the emigration terms *E_ix_*(*t*) and immigration *I_ix_*(*t*) terms. *E_ix_*(*t*) is the number of individuals of species *i* dispersing from patch *x* at time *t*. This is determined by the successful number of 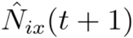 draws from a binomial distribution, each with a probability of *a_i_*. Here, we assume that *a_i_* is equal across all species, except in the scenarios where we include a competition colonization trade-off (see below). Relaxing the assumption of equal dispersal would be a worthwhile next step, but in general we expect that interspecific variation in dispersal would erode the potential for regional scale coexistence unless additional trade-offs are assumed (e.g. competition-colonization trade-off). *I_ix_*(*t*) is the number of individuals of species *i* that arrive via immigration to patch x at time *t* from other patches. We assume that the probability of an individual arriving from another patch decreases exponentially with the geographic distance between patches:

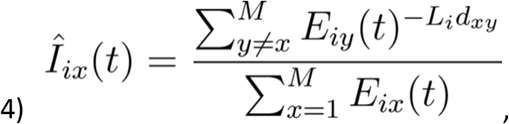

where *Î_ix_*(*t*) is the probability that a dispersing individual of species *i* immigrates to patch *x* at time *t*. *d_xy_* is the geographic distance between patches x and y, and *L_i_* is the strength of the exponential decrease in dispersal with distance *I_ix_*(*t*).

#### Stochasticity

Stochasticity is incorporated in two ways. First, through the stochastic component of immigration and emigration as noted above. Second, we determine the realized population size in the next time step before dispersal 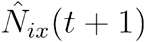 by drawing from a Poisson distribution (following Shoemaker & Melbourne 2016).

### Simulation details

The results presented are based on model simulations of *M* = 100 patch metacommunities with a starting total species richness of *S* = 50 species and an *r_max_* of 5 for all species.

#### Metacommunity spatial structure

We assume that habitat patches are distributed randomly in geographic space with their coordinates drawn from uniform distributions with the range [1, 100] (following Fournier *et al*. 2017; Thompson *et al*. 2017)(Figure S1). We convert these coordinates into a torus to avoid edge effects. Although we chose to restrict our simulations to this one metacommunity size and structure, the model can be run using any number of patches and any spatial structure. Our qualitative results appear robust to variation in species number and metacommunity size, based on initial sensitivity analyses (results not shown) and the fact that portions of our results are qualitatively consistent with other models (e.g. Hubbell 2001; Loreau *et al*. 2003; Mouquet & Loreau 2003; Gravel *et al*. 2006).

#### Density-independent environmental heterogeneity

We assume that the density-independent abiotic environment in a given patch varies continuously between 0 and 1 and is both spatially and temporally autocorrelated (Figure S1, S2). We generate this environmental heterogeneity with an exponential covariance model, using the *RMexp* function in the *RandomFields R* package (Schlather *et al*. 2015). We assume a mean environmental value across time and space of 0.5, an environmental variance of 0.5, and spatial and temporal scales of environmental autocorrelation of 50 and 500, respectively. The model can be easily modified to incorporate other patterns of changes of environmental conditions in time and space.

#### Initialization

We initialize the simulation by seeding each habitat patch with populations of each species drawn from a Poisson distribution where λ = 0.5. Thus, species start in a random subset of patches and at different abundances. We repeat this seeding procedure every 10 time steps over the first 100 time steps, giving each species the opportunity to establish if they can increase from low abundance. Their ability to do so will depend on whether they are suited to the local environmental conditions and their interactions with other species. By seeding the metacommunity randomly, we allow for the possibility of priority effects (but only if the structure of local competition allows for this, otherwise communities will converge in the same environmental conditions). To allow communities to reach equilibrium initially, we hold the environmental conditions in each patch constant for the first 200 time steps, while allowing for spatial variation in conditions.

#### Simulation runs

We ran each simulation for a total of 2200 time steps. This included the 200 time step initialization and an additional 800 time step burn-in period. This duration contained sufficient temporal variation in environmental conditions and community composition to capture dynamics that were representative of particular parameters in the simulation. For each randomly-generated landscape structure, we contrasted a range of dispersal rates *a_i_*, crossed factorially with a range of abiotic niche breadths *σ_i_* and four different structures of competitive effects. To cover the full range, from effectively disconnected to highly connected metacommunities, we varied 15 rates of dispersal, equally distributed in log space from 0.0001 to 0.464. We varied 13 values of niche breadth, equally distributed in log space from 0.001 to 10. These values were chosen to cover the range from so narrow as to preclude species persistence in variable environments to so broad that they are effectively neutral over the range of conditions experienced in the metacommunity. For simplicity, and to make fitness differences solely dependent on fundamental niche match, we assume that all species have the same strength of intraspecific competition *α_ii_* = 1. This is, however, not a necessary condition for generating the dynamics in our model, with the exception of the purely neutral case. The four different structures of density-dependent competitive effects were:

1. equal—*α_ij_* = *α_ii_*;
2. stabilizing—*α_ij_* < *α_ii_* and *α_ij_* is drawn from a uniform distribution in the range [0, 0.5];
3. mixed—values of *α_ij_* are drawn from a uniform distribution in the range [0, 1.5], resulting in a combination of species pairs for which competition is stabilizing, destabilizing, or where one of the two is competitively dominant;
4. competition-colonization trade-off—values of *α_ij_* are drawn from a uniform distribution in the range [0, 1], except for 30% of species, which are considered dominant species. For the dominant species, *α_ij_* > *α_ii_*, and these values are drawn from a uniform distribution in the range [1, 1.5]. Thus, local coexistence is possible among subdominant species, but not between subdominant and dominant species. Coexistence between dominants and subdominants occurs at the regional scale, via the classic competition-colonization trade-off (Hastings 1980). For this, we assume that dispersal rates *α_i_* are an order of magnitude lower for the competitive dominant species compared to the value used for all other species in the community. We also assume that each population has a probability of 0.002 of stochastic extirpation. As a result, this competitive scenario differs from the consistent and continuous assumptions from all other scenarios. Nevertheless, because the competition-colonization trade-off has an integral place as one of the core archetypes in metacommunity theory (e.g. Leibold *et al*. 2004), we include it here for two reasons: first, to demonstrate how such a trade-off can be incorporated in our framework and model (note that other implementations of such a tradeoff are also possible; e.g. Tilman 1994; Shoemaker & Melbourne 2016; Fournier *et al*. 2017); second, to explore the dynamics that emerge from this trade-off across a range of assumptions about abiotic niche responses and dispersal rates.

Note, we scaled all competition coefficients by multiplying them by 0.05, to allow for higher equilibrium abundances. For each of the 30 replicate landscapes, we ran all dispersal rates and abiotic niche breadth scenarios on the same four sets of randomly generated competition coefficients.

#### Response variables

All response variables were based on the dynamics in the simulated metacommunities (e.g. Figure S3, S4) after excluding the first 1000 time steps (200 initialization, 800 burn-in) to avoid initial transient dynamics. Time series were subsampled every 20 time steps to make data file sizes manageable. Only parameter combinations for which species persisted in at least 10 of the 30 replicates were included in the final analysis. Table 1 provides an overview of the metacommunity properties that we calculated in each simulation run, which include multiple richness and abundance metrics and their temporal stability. Simulations were performed in Julia (Bezanson *et al*. 2017), and figures were produced using *ggplot2* in *R* (R Development Core Team 2017) – code available here https://doi.org/10.5281/zenodo.3708350. We have also developed an R package that allows users to simulate metacommunity dynamics using our model - https://github.com/plthompson/mcomsimr. Example dynamics for different combinations of parameters can be explored in this Shiny app - https://shiney.zoology.ubc.ca/pthompson/meta_com_shiny/

**Table 1.**
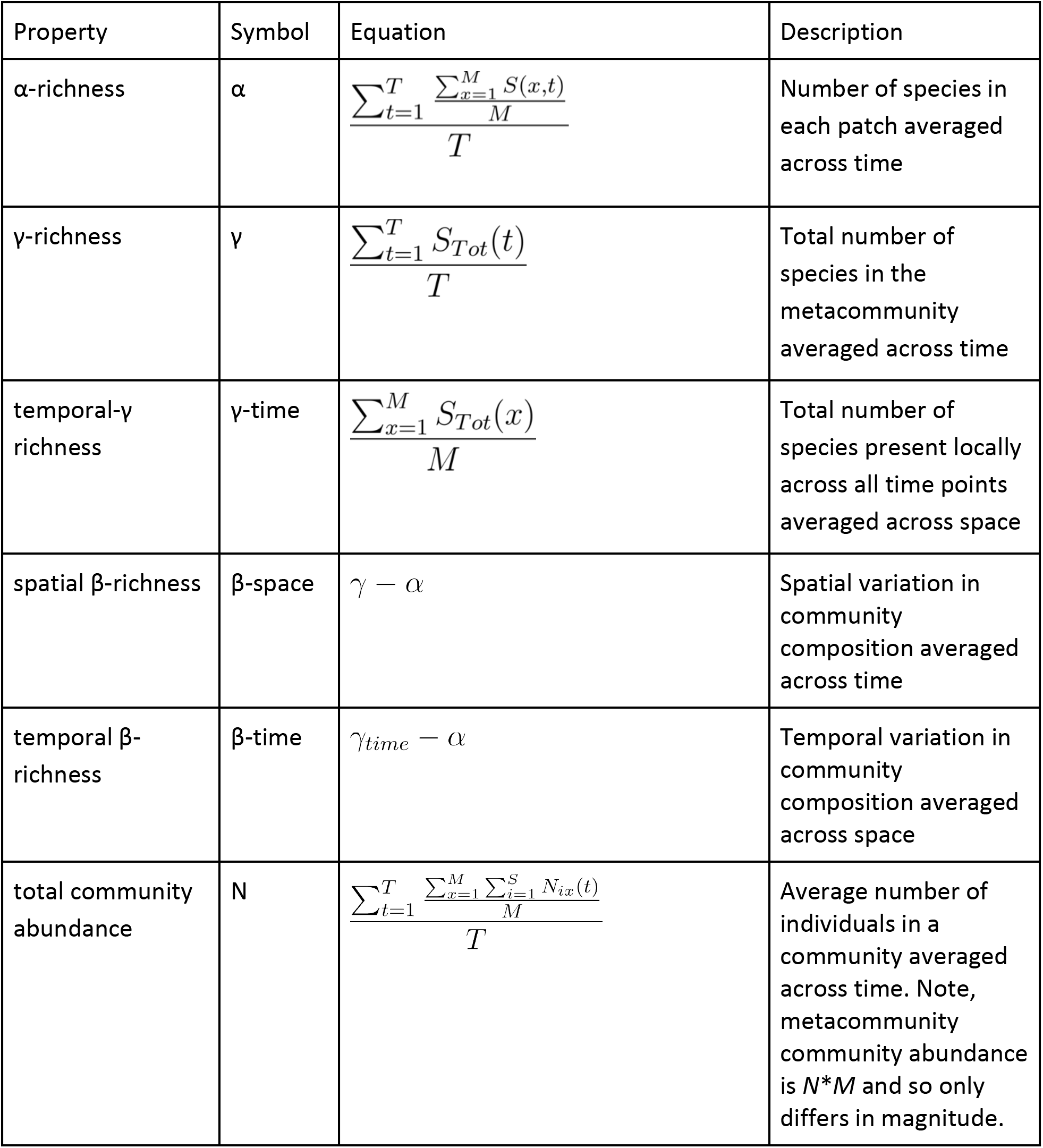
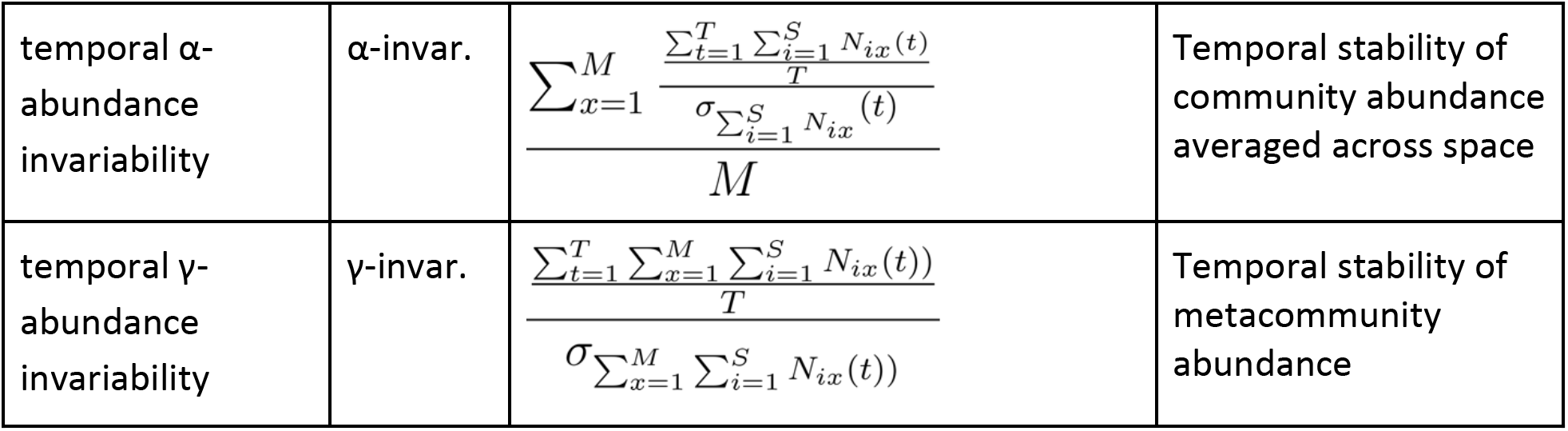
Metacommunity properties calculated from simulated dynamics. *S, M*, and *T* are the number of species, patches, and time points in the simulations, respectively. *S*(*x, t*) is the number of species present in patch *x* at time *t*. *S_Tot_*(*t*) is the number of species present across all patches at time *t*. *S_Tot_*(*x*) is the total number of species present in patch *x* across all time points. σ is standard deviation.

## Results and Discussion

Our overarching message, both within the broader conceptual framework and more specific simulations, is that community dynamics nested within metacommunities exist along a continuum of processes that operate at both local and regional scales. This continuum is defined by three fundamental processes: 1) density-independent responses to abiotic heterogeneity, 2) density-dependent competition, and 3) dispersal—plus stochasticity in how those processes impact community dynamics. The framework generates the dynamics that correspond to classic community and metacommunity ecology theory (e.g. the four metacommunity archetypes; Box 1) but now illustrates how these dynamics change as we vary the strength of the underlying processes. Fully exploring the wide range of dynamics in our model is beyond the scope of a single paper. Instead, we will structure our presentation by discussing the dynamics that result from the three processes and the patterns of diversity, abundance, and stability that they produce. We also highlight key insights from previous community ecology theory, illustrating how these results correspond to specific assumptions about the three processes of our framework.

### The relationship between α, β, and γ-species richness

One classic result regarding how dispersal rates influence local (α-) and regional (γ-) richness, as well as the turnover of species from site to site ( β-richness), comes from Mouquet and Loreau (2003) and Loreau et al. (2003). We can reproduce those predictions (Figure 3a) when we match their assumptions that species have differential and strong responses to abiotic conditions (i.e. narrow abiotic niches) but compete equally for a common resource (i.e. equal inter and intraspecific competition; e.g. *σ_i_* = 0.5 and equal competition). With low dispersal, communities are effectively isolated so different species establish in each patch (low α-, high spatial β-, high γ-richness) and persist through time (low temporal β-richness). Because competition is equal, local coexistence is not possible in the absence of dispersal. As dispersal increases, spatial β- and γ-richness erode, communities have higher turnover through time (temporal β-richness), and α-richness increases. At higher dispersal rates, the metacommunity homogenizes due to source-sink dynamics, spatial and temporal β-richness erodes, and further loss of γ-richness results in losses in α-richness. A key difference between our model and those of Mouquet and Loreau (2003) and Loreau et al. (2003) is that the spatially-explicit nature of our model makes dispersal less effective at homogenizing the metacommunity and reducing γ-richness because we assume that environmental conditions are spatially autocorrelated. Thus, mass effects tend to only allow individual species to dominate subregions of the metacommunity. Because real metacommunities are spatially explicit, we suggest that it is unlikely that dispersal can lead to full homogenization, except in metacommunities composed of a small number of very well-connected patches. Indeed, this may explain why the decline in γ-richness is rarely observed in experimental studies that manipulate dispersal (Grainger & Gilbert 2016).

**Figure 3.**
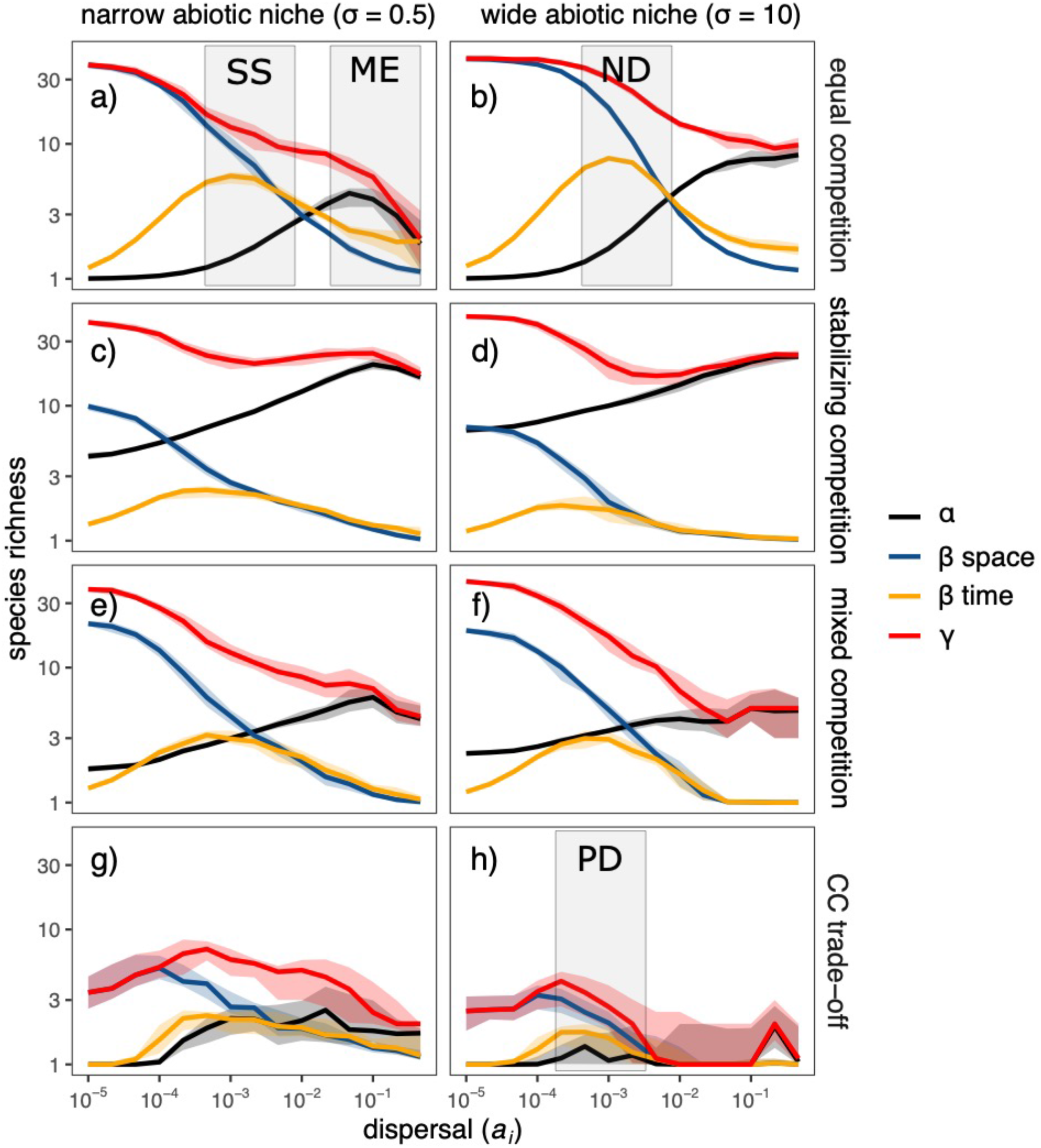
The relationship between dispersal *a_i_* and α-, β- (spatial and temporal) and γ-richness under narrow (column - *σ_i_* = 0.5) and flat (column - *σ_i_* = 10) abiotic niches across the four competitive scenarios (rows). The corresponding parameter space for each of the original metacommunity archetypes: ND - neutral dynamics, PD - patch dynamics, SS - species sorting, and ME - mass effects, is indicated with the shaded boxes. The interquartile range (bands) and median (solid lines) from 30 replicate simulations are shown.

A broadly similar pattern emerges when species abiotic niches are so broad as to be effectively neutral (e.g. *σ_i_*. = 10; Figure 3b). Here again, low dispersal rates result in low α-richness but high spatial β- and γ-richness, but, in this case, as a result of ecological drift, rather than by the match with the abiotic conditions. Just as when abiotic responses are strong, increased dispersal erodes spatial β- and γ-richness but increases α-richness. Temporal β-richness also increases as a result of stochastic colonization and extinction. Of course, like in Hubbell’s (2001) model, this diversity will very slowly decline without speciation or dispersal from outside of the metacommunity.

### Relaxing the assumption of equal competition

Competition in real communities is rarely, if ever, equal, and this can result in a wide range of competitive outcomes (Cushing *et al*. 2004; Ke & Letten 2018). If we assume that interspecific competition is weaker than intraspecific competition, we see that multiple species can coexist locally, even without dispersal (Figure 3c-d). This increased potential for coexistence means that higher rates of dispersal increase α-richness but do not erode γ-richness as much as when competition is equal.

If we instead assume that interspecific competition can be either weaker or stronger than intraspecific competition, we also get species pairs that cannot coexist locally in the absence of mass effects (Ke & Letten 2018). However, because this competitive scenario also includes sets of species that can coexist, multiple community compositions can be present in the same abiotic conditions. Thus, the composition of the community depends on the order in which species arrive and whether they can coexist with species that are already present (i.e. priority effects; Fukami *et al*. 2016). Such priority effects are absent from the traditional archetypes but have been found to influence metacommunity structure (e.g. Shurin *et al*. 2004; Urban & De Meester 2009; Vass & Langenheder 2017; Toju *et al*. 2018). Notably, with this structure of competition, priority effects lead to spatial β-richness, but these are eroded as dispersal homogenizes the metacommunity (Figure 3e, f).

Finally, if we assume that there is a competition-colonization trade-off, subdominant species can only persist by colonizing newly disturbed patches before dominant species arrive. Because of the transient nature of these communities, all scales of diversity are lower and less predictable than in the other scenarios, but follow the same general pattern (Figure 3g,h). Here, we see dynamics that correspond to the classic patch dynamics archetype (Levins & Culver 1971) when we assume intermediate rates of dispersal and flat abiotic niche responses, which corresponds to the common assumption in patch dynamics models that the environment is homogeneous.

### A more general relationship between dispersal and α, β, and γ-diversity

A more comprehensive view can be gained by viewing the relationship with abiotic niche breadth and dispersal at the same time (Figure 4). Here, we see that α-richness is generally highest when dispersal rates are high, abiotic niches are wide, and interspecific competition is weaker than intraspecific competition (Box 2, H1). Spatial β-richness is greatest when dispersal is low (Box 2, H2). Temporal β-richness is greatest when dispersal is intermediate (Box 2, H3). Both spatial and temporal β-richness are promoted by strong competition (e.g. equal or mixed competition) or narrow abiotic niche breadth (Box 2, H4), but high rates of dispersal erode β-richness, regardless of the mechanism responsible for its generation (Box 2, H5).

**Figure 4.**
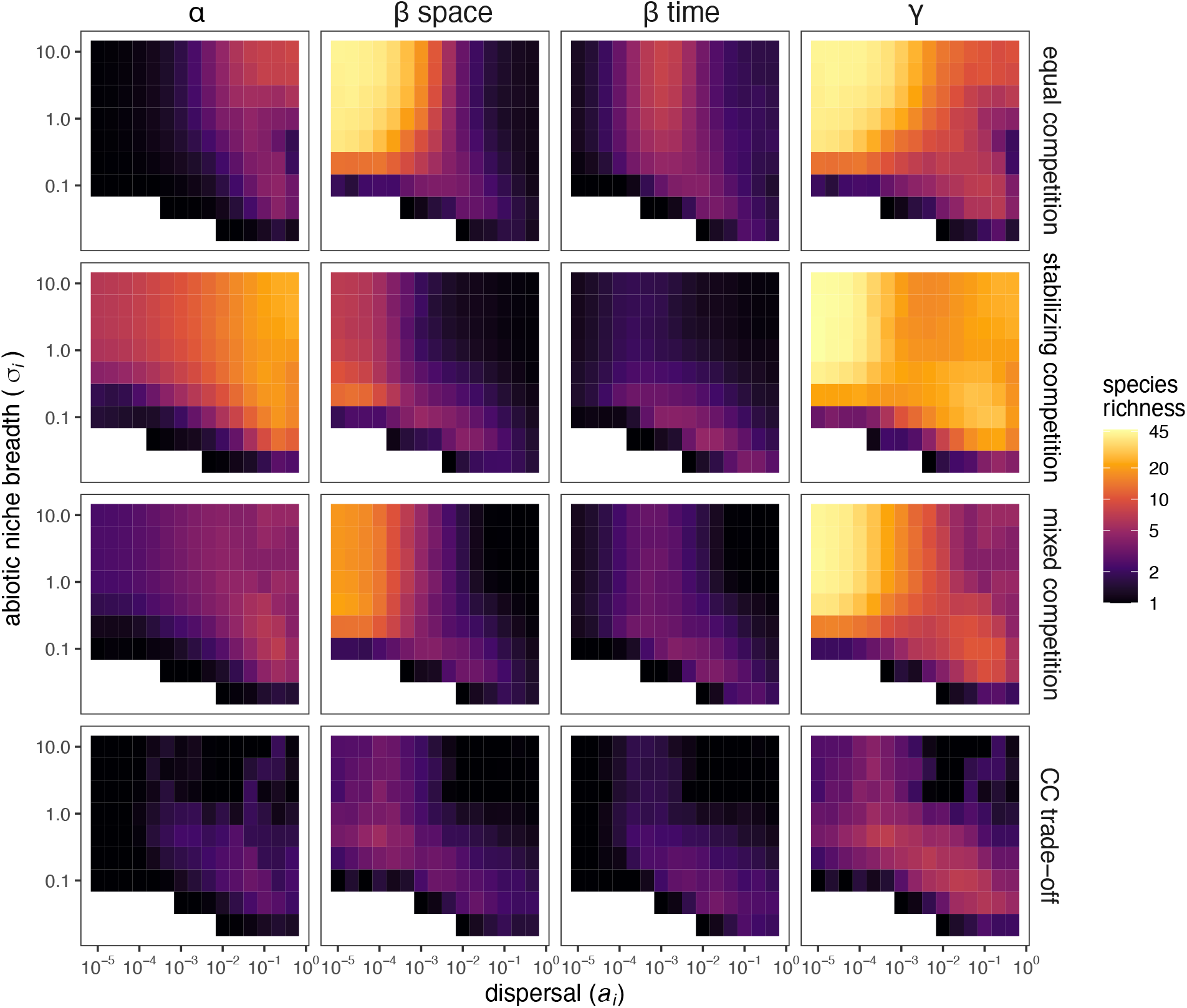
α, β (spatial and temporal), and y richness (columns) across the full range of dispersal rates *a_i_* (x-axis), abiotic niche breadth *σ_i_* (y-axis), and competitive scenarios (rows). Each pixel represents the median value across 30 replicate simulation runs. Colours hues are spaced on a log_10_ scale (see legend). White space represents combinations of parameters where fewer than 10 replicates resulted in species persistence. To see the dynamics that produce these patterns check out our interactive shiny app - https://shiney.zoology.ubc.ca/pthompson/meta_com_shiny/.

#### Box 2 Key hypotheses derived from the model

**H1** – α-diversity (Figure 4, left column) increases with dispersal, as abiotic niche breadths become broader, and as interspecific competition gets weaker (e.g. higher under stabilizing competition compared to equal or mixed competition)
**H2** – spatial beta diversity (Figure 4, 2^nd^ column from left) is highest under low dispersal, regardless of abiotic niche breadths or competition
**H3** – temporal beta diversity (Figure 4, 2^nd^ column from right) is highest under intermediate dispersal, regardless of abiotic niche breadths or competition
**H4** – spatial and temporal β-diversity (Figure 4, middle columns) are both promoted by narrow abiotic niche breadth or strong competition (e.g. equal or mixed vs. stabilizing competition)
**H5** – β-diversity (Figure 4, middle columns) is eroded by high dispersal rates
**H6** –γ-diversity (Figure 4, right column) is eroded by high dispersal, but this occurs more slowly when interspecific competition is weak (e.g. stabilizing competition)
**H7** – metacommunity abundance (Figure 5) becomes less temporally variable as abiotic niche breadth increases or as dispersal increases
**H8** - positive and saturating BEF relationships at local scales (Figure 6) require stabilizing species interactions

Finally, γ-richness is highest with low dispersal and declines as dispersal increases (Box 2, H6). However, γ-richness declines more slowly with dispersal when interspecific competition is weak (Box 2, H6).

### Spatial insurance

The spatial insurance hypothesis predicts that intermediate rates of dispersal should reduce temporal variability in community abundance by allowing species to sort into their preferred habitats as the environment changes (Loreau *et al*. 2003). We find that this effect is strong under the assumptions of Loreau et al. (2003)(Figure 5; equal competition and relatively narrow niche breadth; e.g. *σ_i_* = 0.5), but that it is eroded, lost, or even reversed with other combinations of abiotic niche breadths and biotic interactions. A more general result is that metacommunity abundance becomes more stable (i.e. invariable) with increasing dispersal and with increasing niche breadth (Figure 5; Box 2, H7). A notable exception is that we see a U-shaped relationship between metacommunity invariability and dispersal when some species exhibit destabilizing competition (i.e. mixed competition) and when abiotic niche breadths are nearly flat. This occurs because the strong interspecific interactions result in local compositional states that are relatively unstable and can change dramatically if dispersal introduces a new species that is incompatible with members of the local community.

**Figure 5.**
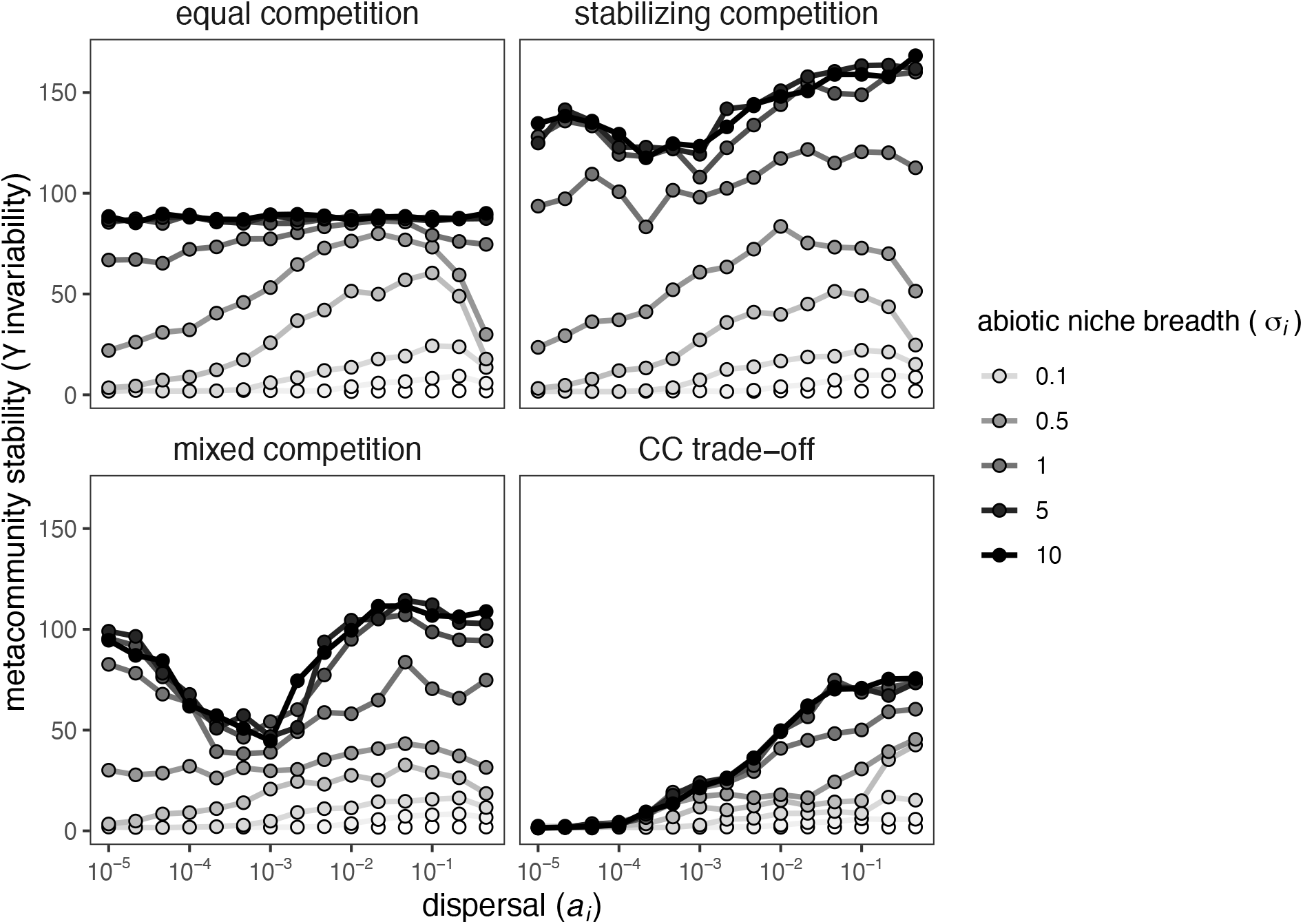
The spatial insurance relationship between dispersal and metacommunity scale stability in total biomass (invariability). Lines connect different dispersal rates *a_i_* with the same abiotic niche breadth *σ_i_* (color of lines and points). Values are medians from 30 replicate simulations. Legend shows the grey scale using a subset of *σ_i_*, values.

### The biodiversity-ecosystem functioning relationship

Understanding how dispersal influences the relationship between biodiversity and ecosystem functioning (BEF) is a key extension of metacommunity theory (Bond & Chase 2002; Loreau *et al*. 2003; Thompson & Gonzalez 2016; Leibold *et al*. 2017). A classic result is that α-richness and community productivity are both maximized at intermediate rates of dispersal (Loreau:2003iz; see also Gonzalez *et al*. 2009; Shanafelt *et al*. 2015; Thompson & Gonzalez 2016). The same set of assumptions reproduce this result in our model when we assume that species are similar in body size and use community abundance as a proxy for productivity (Figure 6; hump-shaped trajectory with equal competition and relatively narrow niche breadth; e.g. *σ_i_* = 0.5). This result, however, depends on abiotic niche breadth and the structure of competition. The BEF relationship at local scales is relatively weak when competition is equal. This is because there is only complementarity in abiotic niche responses, but not resource use, across time and space. Yet, when there is stabilizing competition (Figure. 7), we see the positive and saturating BEF relationships that are common in experimental communities (Box 2, H8) (Cardinale *et al*. 2011; Tilman *et al*. 2014). However, even with stabilizing competition, we see that mass effects at the very highest dispersal rates cause both diversity and functioning to decrease. These BEF relationships remain present, but more variable, under mixed competition and disappear entirely under a competition-colonization trade-off (Figure 6). Together, these findings provide a more comprehensive view of BEF relationships in metacommunities, showing that the strength of these effects can vary dramatically, depending on the underlying mechanisms, and that they are particularly sensitive to how species compete for resources.

**Figure 6.**
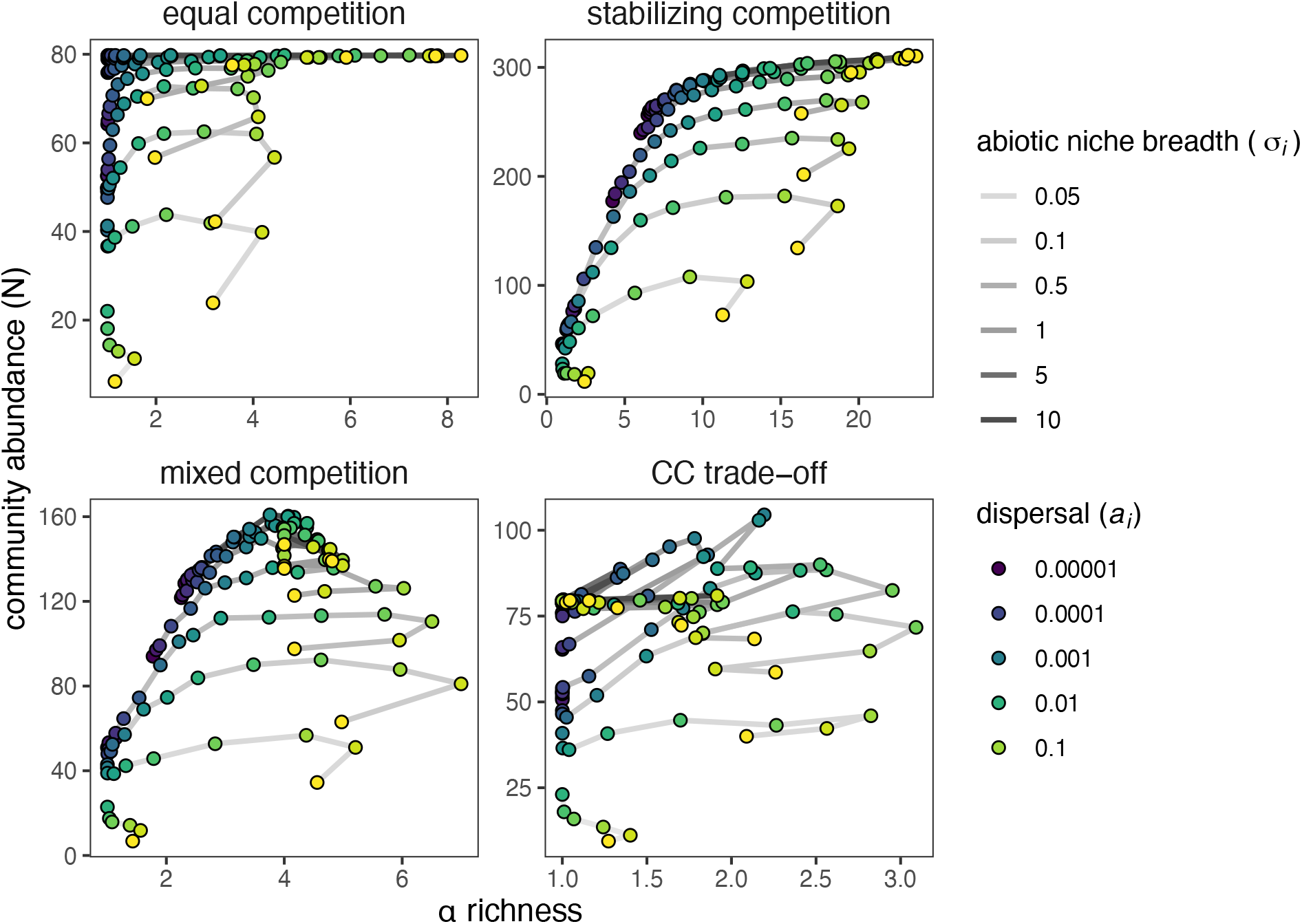
The biodiversity functioning relationship between α-richness and community abundance *N*. Lines connect different dispersal rates *a_i_* (color of point) with the same abiotic niche breadth *σ_i_* (shade of line). Values are medians from 30 replicate simulations. Legend shows the grey scale using a subset of *σ_i_* values and the colour scale using a subset of *a_i_* values.

## Caveats and future directions

Most metacommunity models and theory rely on the assumption that species are equivalent in all or most of their traits (Leibold & Chase 2018). This is most extreme in Hubbell’s neutral theory (2001), but even niche-based models tend to assume that all species have the same dispersal rate, the same shape of the abiotic niche, and usually that competition is equal. Our model breaks one of these assumptions by exploring a range of competitive structures. However, with the exception of the competition-colonization trade-off scenario, we still assume equal dispersal rates and abiotic niche shape within an individual simulation. We did this to demonstrate how changes in dispersal and abiotic niche breadth alter metacommunity dynamics. In nature, however, there is high inter- and intraspecific trait variation, and this can greatly influence community dynamics (Bolnick *et al*. 2011; Violle *et al*. 2012; Roches *et al*. 2017). Exploring the outcome of this variation is an important next step that would allow for the identification of additional trade-offs that may be important for maintaining diversity when species differ in the way they respond to the environment, each other, and space.

In incorporating theory on local biotic interactions and coexistence, we have established clear links to modern coexistence theory (Chesson 2000b). While, our focus has been on how the three-core underlying processes affect the dynamics and diversity of metacommunities, an alternative approach would be to ask how the processes affect the underlying coexistence mechanisms. This would build upon the work of Shoemaker and Melbourne (2016), who quantified the strength of spatial and non-spatial coexistence mechanisms in simulations that corresponded to the classic metacommunity archetypes.

Our focus has been on the ecological dynamics of competitive communities but extending this framework to include trophic dynamics would be a valuable next step. As outlined above, this could be done by including positive and negative *per capita* interaction coefficients (Laska & Wootton 1998; Ives & Cardinale 2004) and different functional responses (Holling 1959). This extension will be critical to connect and synthesize food web ecology (e.g. Holt 2002; Gravel *et al*. 2011; Guzman *et al*. 2019) with the competitive theory that we have focused on here. For example, this would allow for the development of new hypotheses that link the structure of food webs to the underlying metacommunity processes.

Evolution is an additional process that is critical to the dynamics of ecological communities (Hendry 2017) and could be incorporated as a fourth process in the framework. Although beyond the scope of this paper, evolution would affect the underlying traits that determine the three processes that we have focused on. A number of eco-evolutionary metacommunity models have been developed that have focused on the evolution of the density-independent responses of species to abiotic conditions (e.g. de Mazancourt *et al*. 2008; Loeuille & Leibold 2008; Vanoverbeke *et al*. 2015; Leibold *et al*. 2019; Thompson & Fronhofer 2019) or dispersal (e.g. Fronhofer *et al*. 2017). An important next step would be to examine how evolution of density-dependent biotic interactions affects the dynamics of metacommunities. We also did not allow for speciation in our model because it would not have greatly impacted dynamics over the timescales considered. However, it would be important over longer timescales, in particular in parameter combinations that would result in the gradual loss of regional diversity through ecological drift (e.g. Hubbell 2001).

A critical measure of the value of our framework will be in its ability to provide insight into the dynamics of natural communities. We see two ways to achieve this. First, our framework and model provide new hypotheses about how basic ecological processes influence community dynamics (Box 2). These can be tested with observations and experiments to guide data analysis of empirical systems. Second, our model can be used to develop approaches that are informative for identifying the underlying structure of real communities. For example, in a follow-up project, we are using the model with *a priori* specified processes to sample the resulting community time series as a ‘virtual ecologist’ (see also Ovaskainen *et al*. 2019). We can then use these samples of metacommunities with known processes to discern which analytical approaches and metrics are best for distinguishing the underlying processes in natural communities.

## Conclusions

Our framework allows us to recast existing ecological theory in terms of a common set of core processes that underlie the dynamics of communities. This process-based perspective provides a transparent link between local scale coexistence theory and regional scale metacommunity theory. Building on this framework would further advance our understanding of natural communities in a dynamic and changing world, particularly if researchers use it to link other bodies of ecological theory and challenge its predictions through empirical tests in a range of natural systems.

## Acknowledgements

This paper resulted from the sTURN working group funded by sDiv, the Synthesis Centre of the German Centre for Integrative Biodiversity Research (iDiv) Halle-Jena-Leipzig, funded by the German Research Foundation (FZT 118). Additional funding came from Österreichische Forschungsgemeinschaft (ÖFG; International Communication, project 06/15539). We thank all members of the working group for discussions that inspired this manuscript and feedback in its development. We also thank Jacob Usinowicz, Mary O’Connor, and Devin Lyons for valuable discussions and feedback. PLT is supported by Killam and NSERC postdoctoral fellowships. LDM acknowledges KU Leuven Research Fund project C16/2017/002 and FWO project G0B9818. ZH acknowledges support by the Interreg V-A Austria-Hungary program of the European Regional Development Fund (project “Vogelwarte - Madárvárta 2”) and GINOP 2.3.2.-15-2016-00057. LMG. is supported by NSERC CGS-D and UBC Four Year Fellowships. JMC was also supported by the German Centre for Integrative Biodiversity Research (iDiv) Halle-Jena-Leipzig, funded by the German Research Foundation (FZT 118). DSV was supported by the Alexander von Humboldt Foundation and sDiv, the Synthesis Centre of iDiv.

