## Supplemental materials for "A process-based metacommunity framework linking local and regional scale community ecology"

Supplemental Information for **A process-based metacommunity framework linking local and regional scale community ecology**

Patrick L. Thompson, Laura Melissa Guzman, Luc De Meester, Zsófia Horváth, Robert Ptacnik, Bram Vanschoenwinkel, Duarte S Viana, and Jonathan M. Chase

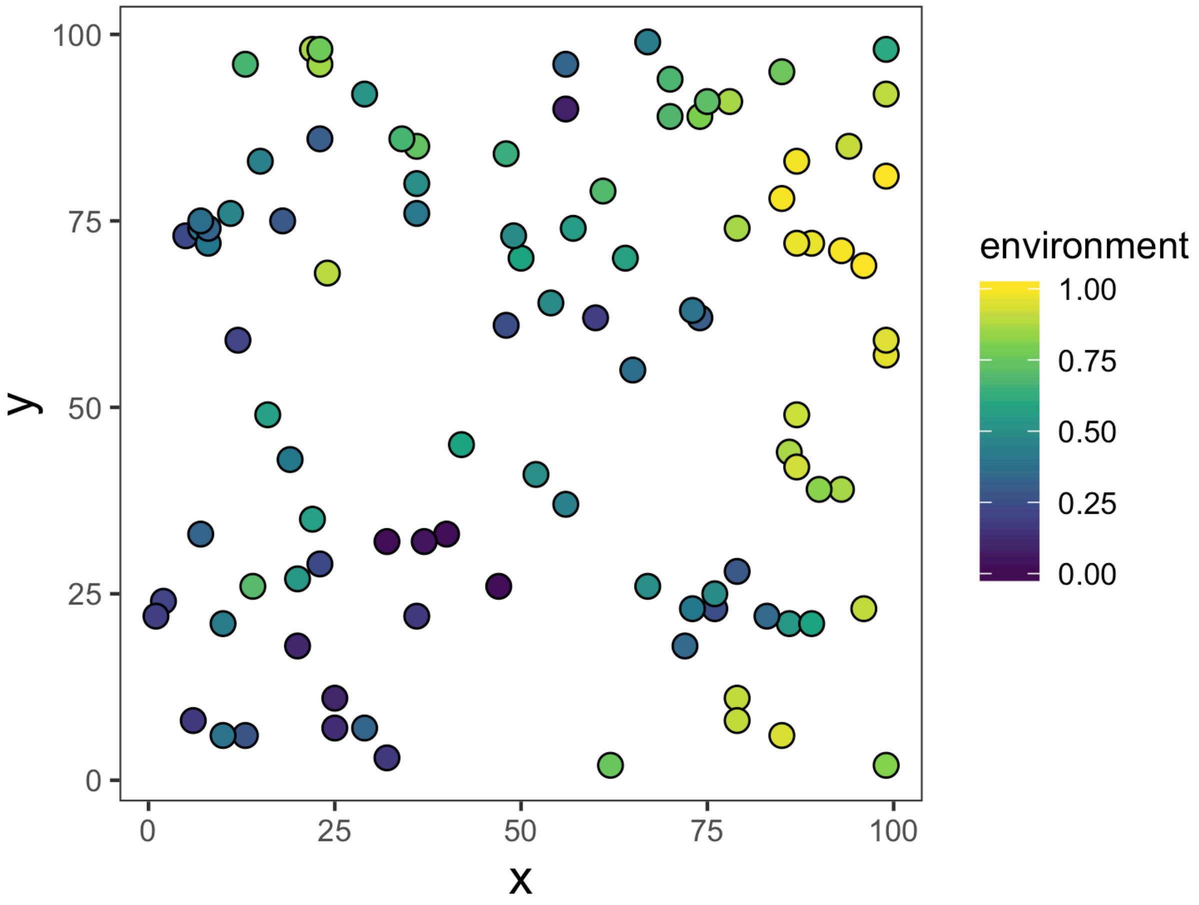


**Figure S1.** Example illustration of one of the 15 replicate landscapes in 2d space (note that the simulation is run by converting this 2d landscape into a torus, which connects the top and bottom as well as the sides of the 2d landscape to avoid edge effects). The patches are shown as dots, and their colour indicates their initial environmental value. Environmental conditions have a spatial autocorrelation in x, y space which is maintained through time as the environment fluctuates.

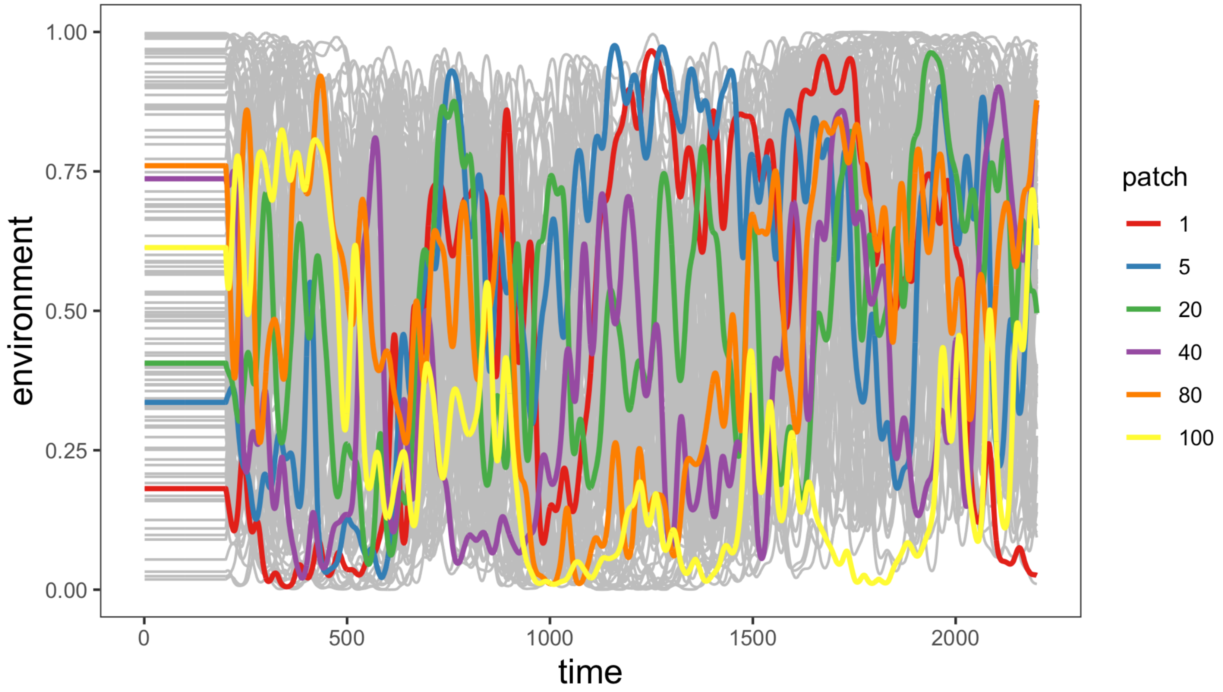


**Figure S2.** Time series of environmental values in the metacommunity from one of the 15 replicate simulations. In each replicate the environment is held constant for the initial 200-time steps to allow the populations to establish. It then varies with a temporal autocorrelation of 500-time steps. The coloured lines highlight the time series for 6 representative patches. The remaining patches are shown as grey lines to show the range of environmental conditions present in the metacommunity at any given point in time.

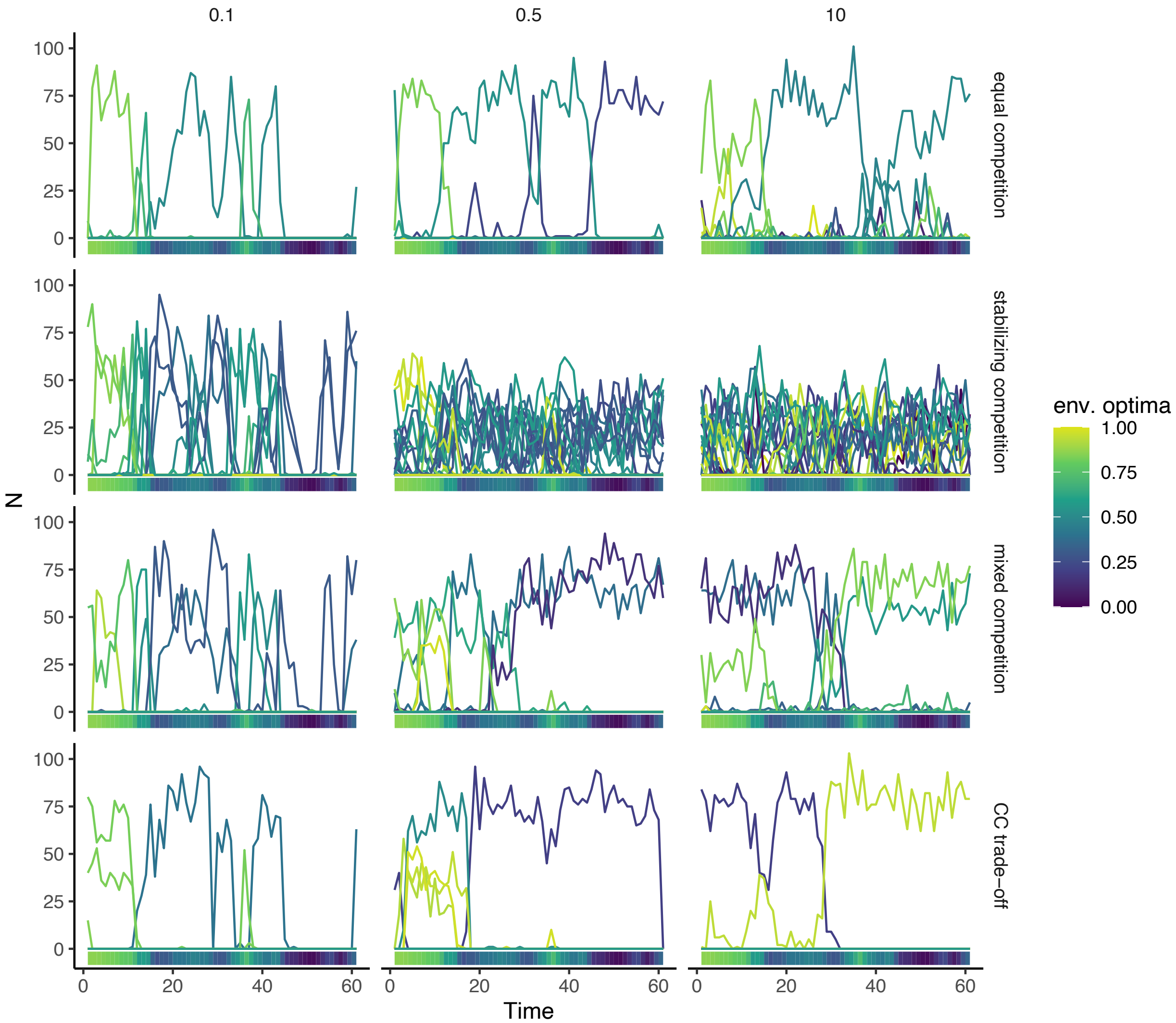


**Figure S3.** Example simulation time series from a single patch in the metacommunity, contrasting three abiotic niche breadths and the four different competition scenarios (rows). Each line in a time series represents the dynamics of a single population, coloured by the environmental optima $z_i$ of that species. The coloured bars at the bottom of each time series show the local environmental conditions and how they change through time. Dispersal $a_i$ is 5e-03 in all cases. To see dynamics across multiple patches and for any combination of parameters check out our interactive shiny app - https://shiney.zoology.ubc.ca/pthompson/meta_com_shiny/.

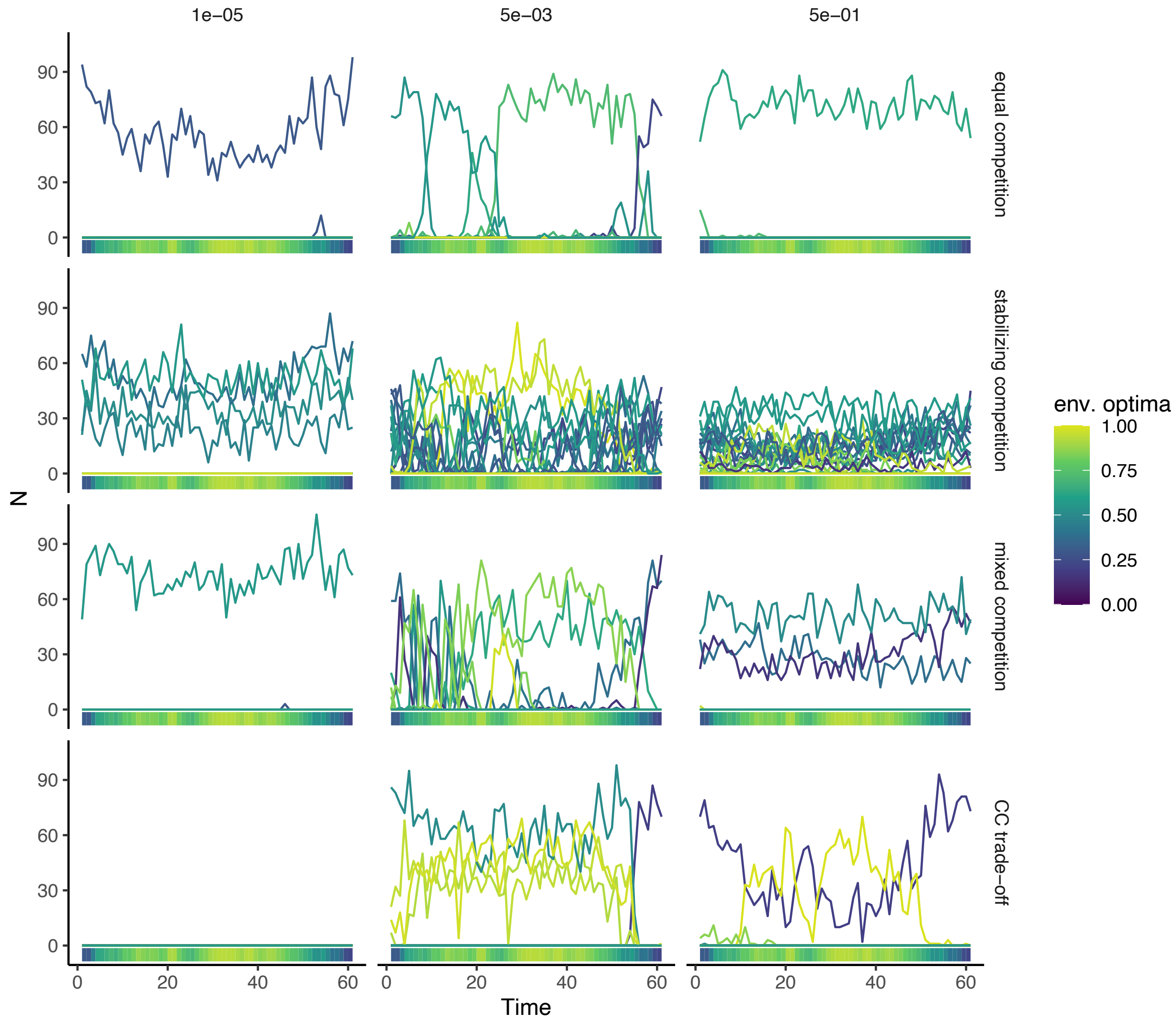


**Figure S4.** Example simulation time series from a single patch in the metacommunity, contrasting three dispersal rates and the four different competition scenarios (rows). Each line in a time series represents the dynamics of a single population, coloured by the environmental optima $z_i$ of that species. The coloured bars at the bottom of each time series show the local environmental conditions and how they change through time. Abiotic niche breadth $\sigma_i$ is 0.5 in all cases. To see dynamics across multiple patches and for any combination of parameters check out our interactive shiny app - https://shiney.zoology.ubc.ca/pthompson/meta_com_shiny/.
